# Mechanistic gene-regulation-based trajectory inference with bowside

**DOI:** 10.64898/2026.10.26.761800

**Authors:** Jan T. Schleicher, Marcello Zago, Manfred Claassen

## Abstract

While single-cell RNA sequencing enables the in-depth study of differentiation processes, simultaneous trajectory inference and gene regulatory network reconstruction remain challenging. We propose bowside, a shallow neural stochastic differentiation model that mechanistically models gene regulation. Our method successfully recovers gene expression trends and the underlying regulatory interactions. We demonstrate bowside on hematopoiesis and induced pluripotent stem cell reprogramming data, retrieving known fate-determining network motifs.

## Main

Cell differentiation involves non-trivial transcriptional changes, regulated by cell-state-specific genetic programs. These changes are induced by transcription factors (TFs) organized in gene regulatory networks (GRNs) and can be studied using single-cell RNA sequencing (scRNA-seq) [1, 2]. Identification of the regulatory targets of these TFs from destructive scRNA-seq snapshot measurements is challenging. While trajectory inference approaches have been used to deduce cell-state sequences [3], the underlying regulatory interactions remain elusive from the resulting trajectories and must be inferred separately using dedicated GRN inference approaches. In contrast to most GRN inference methods, differential equation models stand out by enabling simultaneous trajectory and GRN inference as well as the simulation of GRN behavior, even for unobserved conditions. Stochastic differential equations (SDEs) are particularly well suited for this task as they explicitly model random fluctuations inherent to single-cell data. Neural SDEs provide an efficient model formulation that enables parameter optimization via backpropagation through numerical solutions [4]. However, while deep neural SDEs often achieve a good model fit, they offer limited interpretability and do not allow direct extraction of TF-target interactions.

Here, we introduce bowside, a shallow neural SDE model for GRN-dependent gene expression (Figure 1a). Bowside enables GRN inference, simulation-based trajectory inference, and perturbation prediction. Its input consists of scRNA-seq data along (pseudo)time and an optional prior GRN. The parameters of the SDE, comprising a drift term built around a learnable GRN adjacency matrix and a diffusion term accounting for stochastic fluctuations, are trained by minimizing the discrepancy between numerical simulations and the data at fixed time points. The trained model, with learned SDE parameters, can then address the downstream tasks described above.

**Fig. 1.**
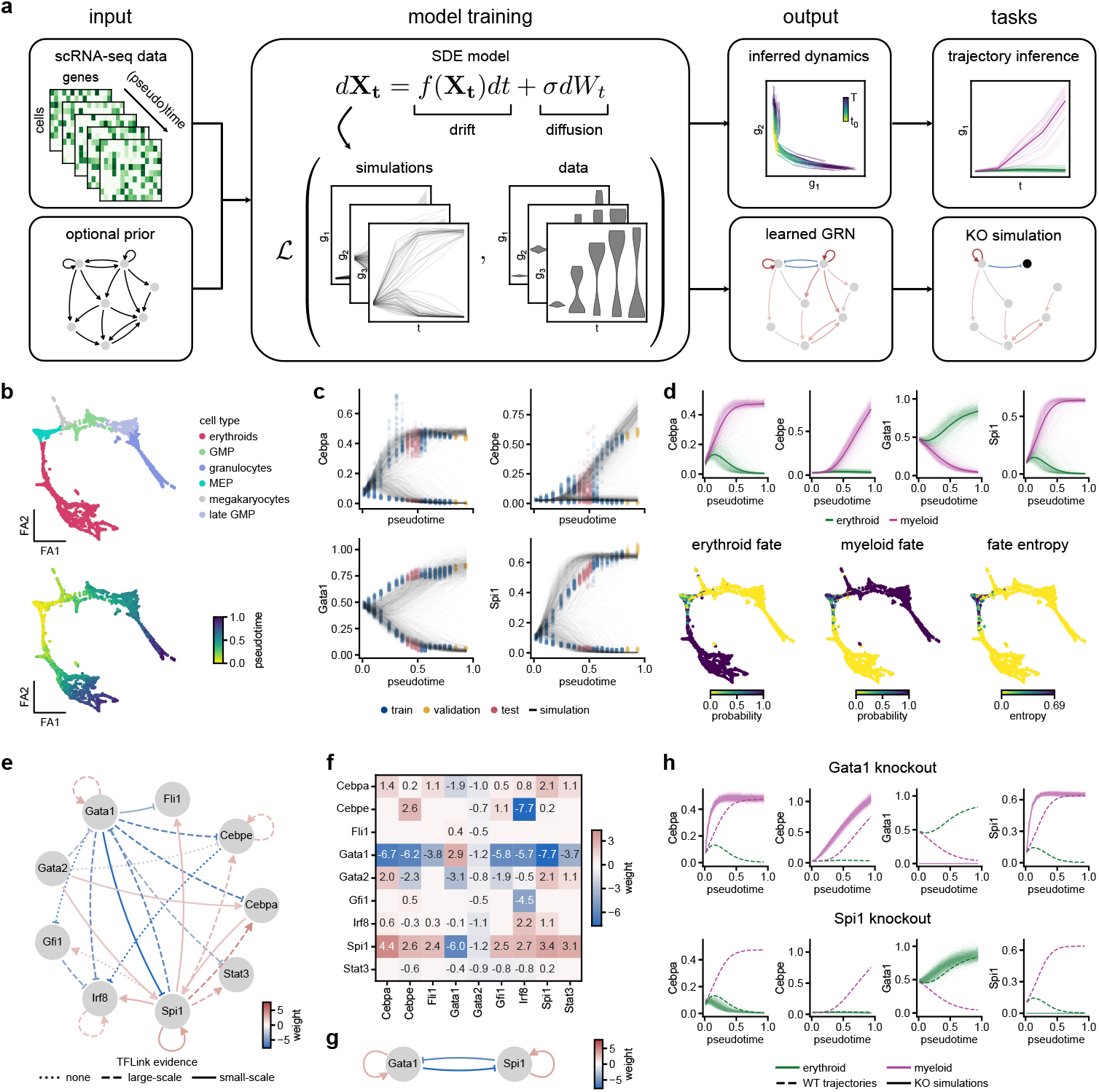
(a) Graphical overview of the bowside SDE model in terms of input, training, output, and downstream tasks. (b) Force-directed graph embedding of hematopoiesis scRNA-seq data colored by cell type (top) and Palantir pseudotime (bottom). (c) Simulations based on the trained model recapitulate branching gene expression. (d) Hierarchical clustering of simulations in gene expression space yields two distinct trajectories (top). Assigning cells in the dataset to the two branches yields fate probabilities (bottom). The entropy over fate probabilities represents differentiation potential. (e) Top 20 edges of the inferred hematopoiesis GRN by absolute weight, highlighting different evidence levels in the TFLink database. (f) Adjacency matrix with annotation of weights *≥* 0.05. Rows: source genes; columns: target genes. (g) Bistable switch between *Gata1* and *Spi1*. (h) Simulation of *Gata1* and *Spi1* knockouts.

We demonstrate bowside on an scRNA-seq dataset of hematopoiesis with Palantir-inferred pseudotime (Figure 1b), choosing nine genes of high relevance (selection criteria: Methods section Hematopoiesis dataset) for this differentiation process [5, 6]. We evaluated the trained model’s dynamics by numerical integration. The simulated expression sequences recapitulated the gene expression data for training, validation, and test time points (Figure 1c, Figure A1a). Based on these simulated sequences, hierarchical clustering recovered two distinct trajectories corresponding to the erythroid and myeloid branches, respectively (Figure 1d, top, Figure A1a). We analyzed cell fate probabilities along pseudotime with a series of Gaussian mixture models on clustered sequences. We inferred high specific fate probabilities for cells in the respective branches and equal fate probabilities for early progenitor cells, coinciding with high differentiation potential quantified by fate-probability entropy (Figure 1d, bottom). We investigated the topology of the learned GRN. Many edges with the highest absolute weight were supported by evidence from the TFLink database [7], and the GRN contained both activating and inhibitory gene regulation (Figure 1e, f). Furthermore, bowside reconstructed a well-known bistable switch between *Gata1* and *Spi1* that determines the erythroid/myeloid fate decision (Figure 1g) [8]. Bowside enables prediction of perturbations of such fate-determining factors. Specifically, simulated knockout of *Gata1* and *Spi1* completely suppressed the corresponding differentiation branches, while simulated knockouts of *Cebpa* and *Cebpe* shifted differentiation towards the erythroid and myeloid branch, respectively, in line with *Cebpa* and *Cebpe* knock-out experiments [5] (Figure 1h, Figure A1b-d), highlighting biologically meaningful knockout simulations.

Additionally, we applied bowside to an scRNA-seq dataset of mouse embryonic fibroblasts (MEFs) undergoing reprogramming to induced pluripotent stem cells (iPSCs) (Figure 2a), selecting 29 TFs highlighted by the original authors [9]. In contrast to the hematopoiesis dataset with pseudotime, this application illustrates that bowside works with experimental time as well. We used 39 time points from day 0 to day 18 of reprogramming. Despite the high sparsity and noise levels, the simulations captured expression trends of iPSC-, trophoblast-, and stromal-cell-related genes (Figure 2b, Figure A2a). For trajectory inference, hierarchical clustering of simulated sequences in gene expression space revealed two to three trajectories with almost equal optimal silhouette scores. The resulting clusters corresponded to the iPSC, trophoblast, and stromal fates (Figure 2c, Figure A2b). Trajectory inference did not capture a separate neural fate described in the original publication, likely because too few cells expressed neural markers at relevant levels (e.g., *Neurod4* in Figure A2a). For the three fates, we observed distinct predetermined cell populations. Some early cells had a high potential to differentiate into any of the lineages (Figure 2d). Finally, the recovered GRN contained activating and inhibitory regulations, with several positive autoregulations (e.g., *Nanog*, *Tdgf1*, *Id3*), feed-forward loops (e.g., between *Id3*, *Prrx1*, and *Nfix*, reinforcing the stromal fate), and mutual inhibition motifs (e.g., *Prrx1* and *Gata3*) (Figure 2e, f).

**Fig. 2.**
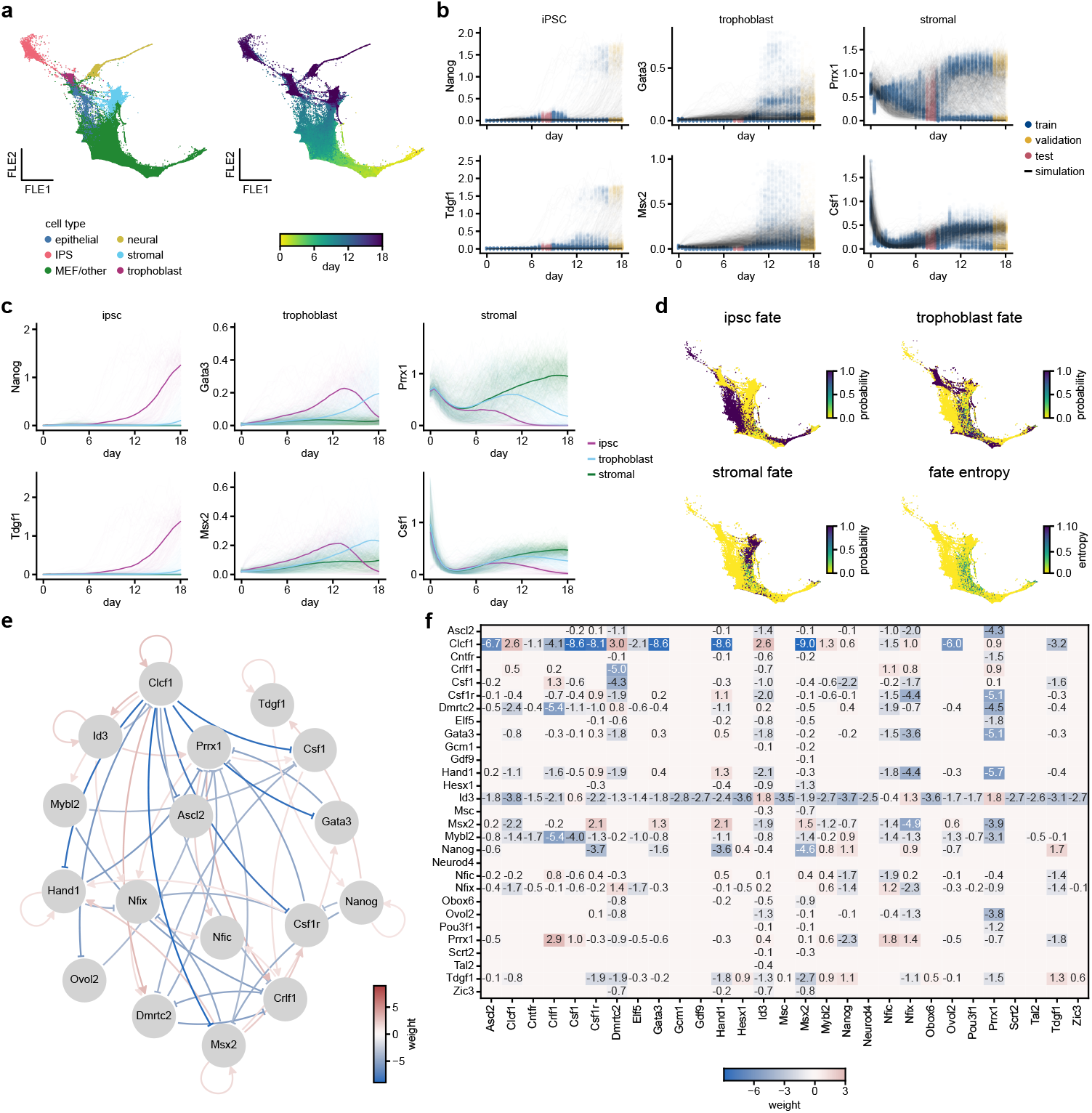
(a) Force-directed layout embedding of iPSC reprogramming scRNA-seq data colored by cell sets (left) and experimental day (right). (b) Simulations based on the trained model recapitulate the expression of iPSC-(left), trophoblast-(center), and stromal-cell-related genes (right) despite high sparsity and noise. (c) Hierarchical clustering of simulations in gene expression space yields three distinct trajectories. (d) Assigning cells in the dataset to the three branches yields fate probabilities. The entropy over fate probabilities represents differentiation potential (bottom right). (e) Top 25 activating and inhibitory edges, respectively, of the inferred GRN by weight, omitting genes that are isolated after selecting only the top edges. (f) Adjacency matrix with annotation of weights *≥* 0.05. Rows: source genes; columns: target genes.

In conclusion, bowside constitutes a mechanistic GRN-based trajectory inference method. Its SDE formulation enables (1) straightforward interpretation by direct extraction of regulatory interactions, as well as efficient (2) trajectory inference and (3) perturbation prediction by simulation. The model recovered expression dynamics and proposed underlying GRNs for scRNA-seq datasets of hematopoiesis and iPSC reprogramming. For both datasets, the inferred GRN weights were stable across multiple repetitions of model fitting (Figure A3). In benchmarks for GRN and cell fate inference on four synthetic datasets, bowside performed competitively with baseline methods (Figure A4). Unlike GRN baseline methods, bowside provides not only a GRN estimate but also a model whose dynamics can be simulated, particularly under new, unobserved conditions. It can be applied to studies providing either pseudotime or real experimental time if enough time points are available. The temporal resolution must be sufficient to capture relevant gene expression changes of the studied process. However, because our training paradigm uses cells sampled at fixed (pseudo)time points, a major limitation is that the differentiation trajectories and corresponding endpoints must be temporally aligned. Individual branches progressing at different speeds violate the model assumptions. This occurs if branch endpoints have strongly different pseudo-times. In this scenario, the model assumes converging trajectories rather than distinct endpoints. To increase the applicability of bowside, future work should explore additional ways to prepare appropriate training data from available scRNA-seq datasets. For instance, training sequences can be generated from a Markov chain process, e.g., based on RNA velocity [10]. In this case, individual sequences could be aligned with dynamic time warping to sample cells from the synchronized sequences at fixed time points.

## Methods

### The bowside model

We model dynamic gene expression using a shallow neural stochastic differential equation (SDE) model based on a gene regulatory network (GRN). The GRN for a set of *g* genes is represented by the weighted adjacency matrix ***A*** ∈ ℝ*^g×g^*, where *A_ji_* is the weight of the edge from gene *j* to gene *i*. The rate of change of the expression *x_i_* of a gene *i* at time *t* follows

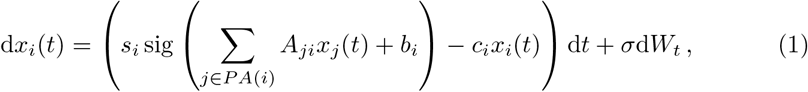

where *s_i_*, *b_i_*, and *c_i_* denote a gene-specific scaling factor, bias, and degradation rate, respectively; sig is the sigmoid activation function; *σ* is the diffusion coefficient; and *W* denotes a Wiener process.

In matrix form, we can express the model as

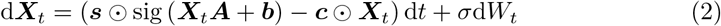

for an expression matrix ***X****_t_* ∈ ℝ*^n×g^* with number of cells *n* and number of genes *g* at time *t*, scaling factors ***s*** ∈ ℝ*^g^*, adjacency matrix ***A*** ∈ ℝ*^g×g^*, biases ***b*** ∈ ℝ*^g^*, and decay rates ***c*** ∈ ℝ*^g^*. These vectors of size *g* are applied element-wise to the rows of the *n* × *g* matrices (⊙ represents element-wise multiplication). The diffusion coefficient *σ* can be either a scalar that is the same for each gene or a vector (***σ*** ∈ ℝ*^g^*) with a distinct value for each gene.

In another variation of the model, the diffusion coefficient is multiplied by the expression matrix to make the noise dependent on the current gene expression state:

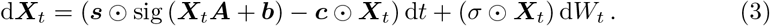

The choice between the constant-noise model in Equation 2 and the multiplicative-noise model in Equation 3 is a hyperparameter that hyperparameter optimization can adjust, as described in subsection Model fitting and hyperparameter optimization.

### Model implementation

We implemented the bowside model in PyTorch [11] as an Itô SDE with diagonal noise to be compatible with the torchsde library [12]. The adjacency matrix is represented as a float tensor of shape *g* × *g*. Additionally, the weighted adjacency matrix is multiplied by a binary mask of the same shape that indicates the presence (1) or absence (0) of edges. Prior knowledge of possible gene regulations can be used to initialize the adjacency matrix and the mask. If only the presence of edges is known, a binary matrix can be used. If information about the direction of regulation is available as well, negative regulations can be encoded by the value −1 in the initialization array. In either case, the weighted adjacency matrix can be initialized with values from the initial (binary or ternary) adjacency matrix or multiplied by values drawn from a normal distribution, preserving the sign of the regulation directions. Without prior information, the mask is initialized as a matrix of ones, representing a fully-connected GRN, and the adjacency matrix is initialized as a sparse matrix with non-zero values drawn from a normal distribution with mean 0 and standard deviation 1.

Scaling factors, biases, and decay rates are represented by their natural logarithm to ensure positivity. Initial log bias and log decay rate values are drawn from a normal distribution with mean 0 and standard deviation 0.01. The initial scaling factors are set to 1. The diffusion coefficient can be fixed (only with a shared value for all genes) or learnable (shared or gene-wise). To avoid negative values when the coefficient is learned, we represent it as its natural logarithm.

### Dataset setup and model training

The bowside model is trained based on scRNA-seq data. Specifically, we use an AnnData object [13] as input for creating a custom PyTorch [11] pseudotime dataset. The prerequisite is the presence of (pseudo)time values in the cell-level annotations of the AnnData object. The cells are then binned based on their (pseudo)time. For real, experimental time, the true discrete time points can be used as the bins. For pseudotime, we instead form bins based on pseudotime quantiles.

To generate the pseudotime dataset, we sample a fixed number of cells for each time point with replacement. Using the gene expression of the cells in the respective bins, we obtain a series of multidimensional distributions of expression vectors at each time point.

We train the model by gradient descent with the RMSprop [14] or Adam optimizer [15]. In each training epoch, the cells at the first time point are used as the initial values for numerical integration of the SDE. The resulting simulations, based on the current parameters of the model, are compared to the pseudotime data with a distribution-based loss function (see below for details). Then, the parameters (weighted adjacency matrix, scaling factors, biases, decay rates, and diffusion coefficients) are adjusted by computing the gradient of the loss function by backpropagation through the SDE solver and a step of the optimizer. No gradient is computed for the binary mask of the adjacency matrix. These two steps are done either for the whole dataset at once or iteratively for multiple batches of cells.

In addition to optimizing the model parameters, we implemented a pruning step for the adjacency matrix, executed every *n*-th epoch and at the end of training. Here, any edges with a weight close to 0 (with specified absolute and relative tolerance) are removed from the model by setting the corresponding entry of the mask to 0.

### Loss function

We use the Sinkhorn divergence, an unbiased, entropically regularized Wasserstein distance, as the loss function between true and simulated cell populations at each time point [16]. The Wasserstein distance, also known as the optimal transport or earth mover’s distance, measures the minimum cost of converting one probability distribution into another. Given two probability measures *α* and *β* on X ⊆ ℝ*^d^* and a cost function *c*: X × X → ℝ*_≥_*_0_, their Wasserstein distance is

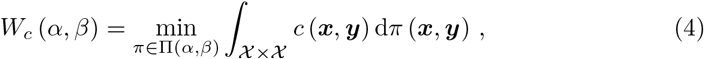

where the minimization is over all possible coupling measures Π(*α, β*), i.e., probability measures on ***X*** × ***X*** whose marginals are *α* and *β*. In this work, we use the squared Euclidean distance as the cost function:

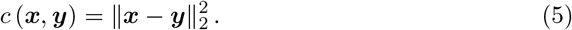

The entropically regularized Wasserstein distance, a computationally more efficient approximation of optimal transport cost [16], is defined as

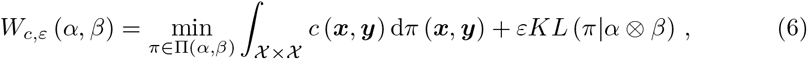

where 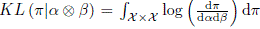 is the Kullback-Leibler (KL) divergence of the coupling *π* from the product measure of *α* and *β*, and *ε >* 0 is the strength of the entropic regularization. However, the regularization introduces bias, so *W_c,ε_* (*α, α*) ≠ 0. The Sinkhorn divergence corrects for this bias and is defined as

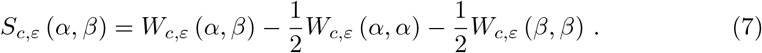

For our implementation of the Sinkhorn divergence loss function, we used the GeomLoss package [16]. We further extended the loss to use different weights for the individual time points and to add an L1 regularization on the weights of the adjacency matrix. Accordingly, for true gene expression ***X*** and predicted expression 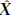 at time points *t* = 0, …, *T* , we compute the loss as

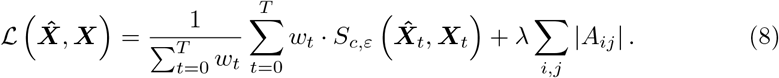

### Trajectory inference

After training the model as described above, gene expression over time is simulated by numerical integration of the SDE system from the initial time point. The system is evaluated at the *n_t_* time points in the dataset to generate *n* sequences of the expression of *g* genes spanning the entire process, represented by an array 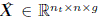. Trajectories in gene expression space are inferred through hierarchical clustering of the simulated sequences. To this end, the temporal dimension is flattened by concatenating the individual time points *t* = 0*, . . . , T* to obtain an array 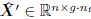:

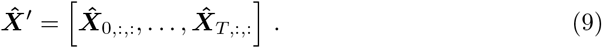

Then, pairwise Euclidean distances between the (*g* · *n_t_*)-dimensional sequences are computed, and a hierarchical tree is constructed with Ward linkage. The tree is cut at different depths to obtain *k* trajectories, and the optimal number of trajectories, *k^∗^*, is obtained by computing the silhouette coefficients for different clusterings and choosing the clustering with the highest value.

To compute fate probabilities for the cells in the original dataset with respect to the inferred trajectories, we construct a Gaussian mixture model for each time point. For this, we calculate the mean and covariance for each cluster *c* over the corresponding sequences. Based on the cluster labels *y_i_* ∈ [1*, . . . , k^∗^*], the index set of all sequences belong to cluster *c* is defined as

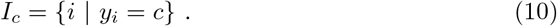

Then, the cluster mean vector ***µ****_c,t_* ∈ ℝ*^g^* and covariance matrix **Σ*_c,t_*** ∈ ℝ*^g×g^* at time point *t* are computed as

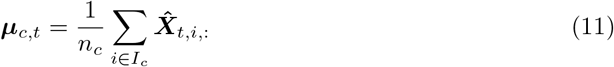

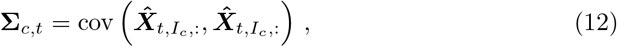

where cov is the unbiased sample covariance, with the columns of 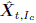 representing the variables (i.e., genes), and its rows representing the samples. This yields a *g*-dimensional Gaussian mixture model for each time point, where the overall probability of a cell ***x****_t_* ∈ ℝ*^g^* corresponding to time point *t* being drawn from the mixture model is defined as

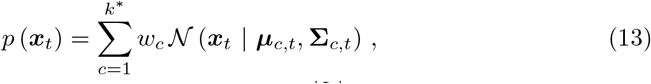

with the individual mixture component weights 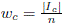. Accordingly, the fate probability of a cell ***x****_t_* with respect to the fate, i.e., trajectory, *c*, is the responsibility of the corresponding component

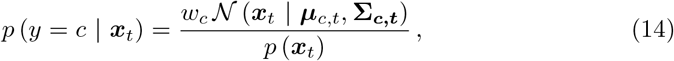

where *y* is the cell’s trajectory assignment.

The differentiation potential of a cell is represented by the entropy over its fate probabilities for all trajectories *c* ∈ [1*, . . . , k^∗^*],

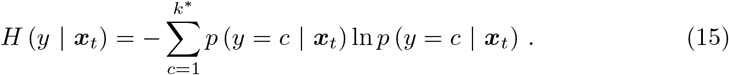

The maximum differentiation potential of a cell is obtained if all trajectories are equally likely, i.e., 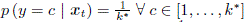. Accordingly, the corresponding maximum value is 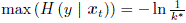.

### Knockout simulation

To simulate the knockout of a gene *i*, we set its expression to zero and remove all its outgoing edges from the GRN before numerical integration by setting the corresponding values in the adjacency matrix to zero, i.e., ***A****_i_*,: ← **0**. All other parameters retain their learned values. We obtain knockout simulations 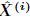 by numerically integrating the adapted model and evaluating the system at the *n_t_* time points in the dataset.

### Datasets and preprocessing

All datasets were obtained in AnnData format [13] and processed with scanpy (v1.11.5) [17] and Palantir (v1.4.1) [6] in Python.

#### Hematopoiesis dataset

A hematopoiesis scRNA-seq dataset by Paul *et al.* [5] containing the expression matrix, cell type annotations, diffusion pseudotime estimates [18], a principal component analysis (PCA) projection, and a force-directed graph embedding used for visualization was downloaded through the CellOracle package with celloracle.data.load Paul2015 data() [19]. The normalized, unlogarithmized counts stored in the layer “raw count” of the AnnData object were log-transformed with scanpy.pp.log1p.

We used Palantir to impute the log-transformed counts with MAGIC [6, 20]. To this end, diffusion maps were computed based on the PCA projection with palantir.utils.run diffusion maps with five diffusion components and otherwise default parameters (i.e., 30 nearest neighbors). Then, the multiscale space was determined with palantir.utils.determine multiscale space, and counts were imputed with palantir.utils.run magic imputation, using default parameters.

For pseudotime estimation with Palantir, the pre-annotated root cell in adata.uns[“iroot”] was used as the start cell, and the cells with the highest diffusion pseudotime in the Erythroids, Monocytes, and Granulocytes cell types, respectively, were set as the terminal states. Palantir was run with 500 waypoints and otherwise default parameters using the function palantir.core.run palantir. For further analysis, we removed monocytes from the dataset because their endpoint pseudotime was lower than that of the other two terminal states. Based on multiple publications, we annotated the following nine TF genes as hematopoiesis-relevant genes to use for model fitting: *Cebpa*, *Cebpe*, *Fli1*, *Gata1*, *Gata2*, *Gfi1*, *Irf8*, *Spi1*, and *Stat3* [6, 19, 21, 22]. Other potentially relevant genes described in the respective publications were excluded if they were not present in the dataset.

#### iPSC reprogramming dataset

An scRNA-seq dataset of mouse embryonic fibroblasts (MEFs) undergoing reprogramming into induced pluripotent stem cells (iPSCs) from Schiebinger *et al.* was downloaded from the Waddington optimal transport package tutorial at https://broadinstitute.github.io/wot/tutorial/ [9]. Specifically, we obtained the expression matrix with log-transformed counts per 10,000 in H5AD format, along with additional text files containing metadata on force-directed layout embedding coordinates, experimental time points, cell sets, and experimental condition (serum or 2i treatment). We subsetted the dataset to the serum condition for further analysis.

The log-transformed counts were scaled, and PCA was performed with default parameters. Based on the PCA projection, diffusion maps were computed with ten diffusion components, using 200 nearest neighbors because of the large dataset size with palantir.utils.run diffusion maps. The multiscale space was computed with palantir.utils.determine multiscale space, and MAGIC imputation of log-transformed, unscaled counts was performed using palantir.utils.run magic imputation with default parameters.

Reprogramming genes were annotated based on Figure 7 of the original publication [9]. We further grouped the genes by cell set (mesenchymal-epithelial transition (MET), stromal, epithelial, trophoblast, neural, iPSC) based on the same figure (Table 1).

**Table 1.** Selected reprogramming genes by cell set.

| Cell Set | Genes |
| --- | --- |
| MET | *Dmrtc2, Pou3f1, Zic3* |
| stromal | *Clcf1, Crlf1, Csf1, Gdf9, Id3, Nfic, Nfix, Prrx1* |
| epithelial | *Ascl2, Elf5, Gata3, Ovol2* |
| trophoblast | *Csf1r, Gcm1, Hand1, Msx2* |
| neural | *Cntfr, Neurod4, Scrt2, Tal2* |
| iPSC | *Hesx1, Msc, Mybl2, Nanog, Obox6, Tdgf1* |

### Model fitting and hyperparameter optimization

For model fitting and hyperparameter optimization, we constructed custom PyTorch pseudotime datasets from the scRNA-seq datasets, as described in subsection Dataset setup and model training. For the hematopoiesis dataset, we binned cells into 20 quantiles along Palantir pseudotime. For the iPSC dataset, we used the 39 experimental time points, ranging from day 0 to day 18 in half-day increments, with two additional time points at day 8.25 and day 8.75. For both datasets, we sampled 1,000 cells per time point with replacement, using the MAGIC-imputed expression data of the selected genes.

To obtain separate training, validation, and test sets, we split the datasets by time point. Based on fixed proportions of time points in the validation and test sets, *r_val_* and *r_test_*, the last ⌊*r_val_* · *n_t_*⌋ time points were used as the validation dataset for early stopping and hyperparameter tuning, and the middle ⌊*r_test_* · *n_t_*⌋ time points of the remaining time points were set aside for interpolation testing. We used all other time points to train the model parameters.

The weights *w_t_*of the loss function in Equation 8 were generated to range from 0 for the first to 10 for the last time point, following a square root function, to emphasize later time points more strongly during training:

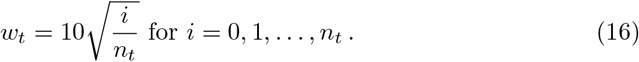

We split the resulting weights into training, validation, and test weights according to the corresponding time-point sets.

Hyperparameters were tuned with Optuna [23]. We used the default TPESampler to efficiently explore the space of both continuous and categorical hyperparameters, running 200 trials per dataset. For each trial, a model was trained with the current sampled hyperparameter values, using only the training time points to optimize the SDE parameters. We used early stopping with a patience of 100 epochs based on the validation loss and treated the final validation loss of each trial as the objective of the hyperparameter optimization. We chose the model with the lowest validation loss after training for further evaluation. Table 2 shows the fixed hyperparameter values and the tested ranges or choices for the tuned hyperparameters. The optimal hyperparameters are shown in Table 3.

**Table 2.** Model training hyperparameters. Square brackets indicate a range of tested values, either sampled on a linear or logarithmic scale, and curly brackets indicate a categorical choice.

| Group | Hyperparameter | Value |  | Sampling |
| --- | --- | --- | --- | --- |
|  |  | Hematopoiesis | Reprogramming |  |
| integration | dt | 0.01 | 0.1 | n.a. |
|  | solver | {Euler, Milstein} |  | categorical |
| data split | train proportion | 0.7 | 0.8 | n.a. |
|  | test proportion | 0.2 | 0.1 | n.a. |
|  | validation proportion | 0.1 | 0.1 | n.a. |
| training | L1 reg. strength | $[10^{-5}, 10^{-2}]$ | | logarithmic |
| | learning rate | $[10^{-5}, 10^{-1}]$ | | logarithmic |
|  | optimizer | {Adam, RMSprop} |  | categorical |
|  | epochs | [500, 5000] |  | linear |
|  | patience | 100 |  | n.a. |
| model setup | normal dist. adjacency init | {True, False} |  | categorical |
| | $\sigma$ | $[10^{-2}, 10]$ | | logarithmic |
| | shared $\sigma$ | {True, False} | | categorical |
| | fixed $\sigma$ | {True, False} | | categorical |
|  | non-multiplicative noise | {True, False} |  | categorical |

**Table 3.**
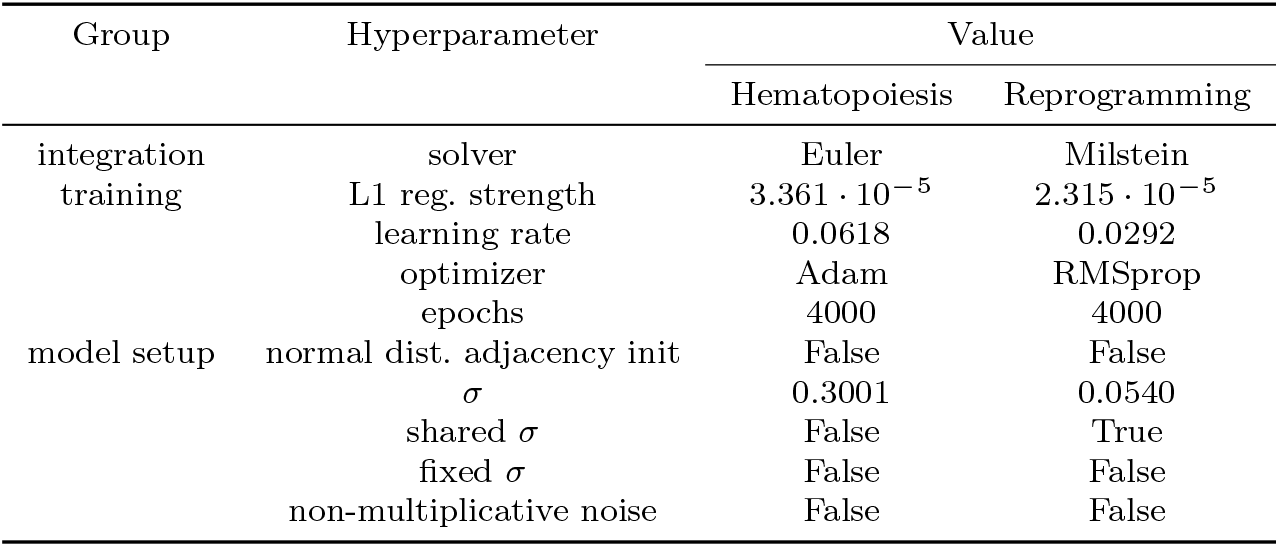
Optimal hyperparameters.

To investigate the stability of the inferred GRN weights, we trained ten separate models per dataset with the respective optimal hyperparameters. Then, we computed the mean and standard deviation of the adjacency matrices across the ten separate models. We used Wilcoxon signed-rank tests with Benjamini-Hochberg false discovery rate (FDR) correction to determine whether the inferred weights for each possible edge were significantly different from 0. Additionally, we computed Spearman’s rank correlation for all pairwise combinations of the individual models’ weighted adjacency matrices to asses the stability of the edge weight ranking.

### Benchmarking GRN inference

We used synthetic datasets from BEELINE based on four curated GRN models to benchmark GRN inference [24]. For each of these four curated networks, we downloaded ten distinct simulated datasets hosted on Zenodo at https://doi.org/10.5281/zenodo.19009603 [25] and converted them to AnnData objects [13]. To avoid conflating GRN inference performance with imputation efficiency, we used the dataset versions without simulated dropout. For all datasets, the same preprocessing steps were performed using scanpy (v1.11.5) [17]. The expression matrices were scaled per gene. We computed neighborhood graphs with 30 nearest neighbors based on scaled gene expression. Because the cell-level metadata in the downloaded datasets contained separate pseudotimes for different trajectories, and each cell belonged to only one trajectory, we created a combined pseudotime annotation comprising all trajectory-level pseudotimes.

We fitted the model and optimized hyperparameters as described in subsection Model fitting and hyperparameter optimization, binning cells into 20 pseudotime quantiles and sampling 200 cells per quantile, using only the first of the ten datasets for each curated network. Table 4 shows the hyperparameter values tested in the Optuna optimization [23]. The optimal hyperparameters were used to train ten separate models per curated network, one for each of the ten distinct synthetic datasets.

**Table 4.**
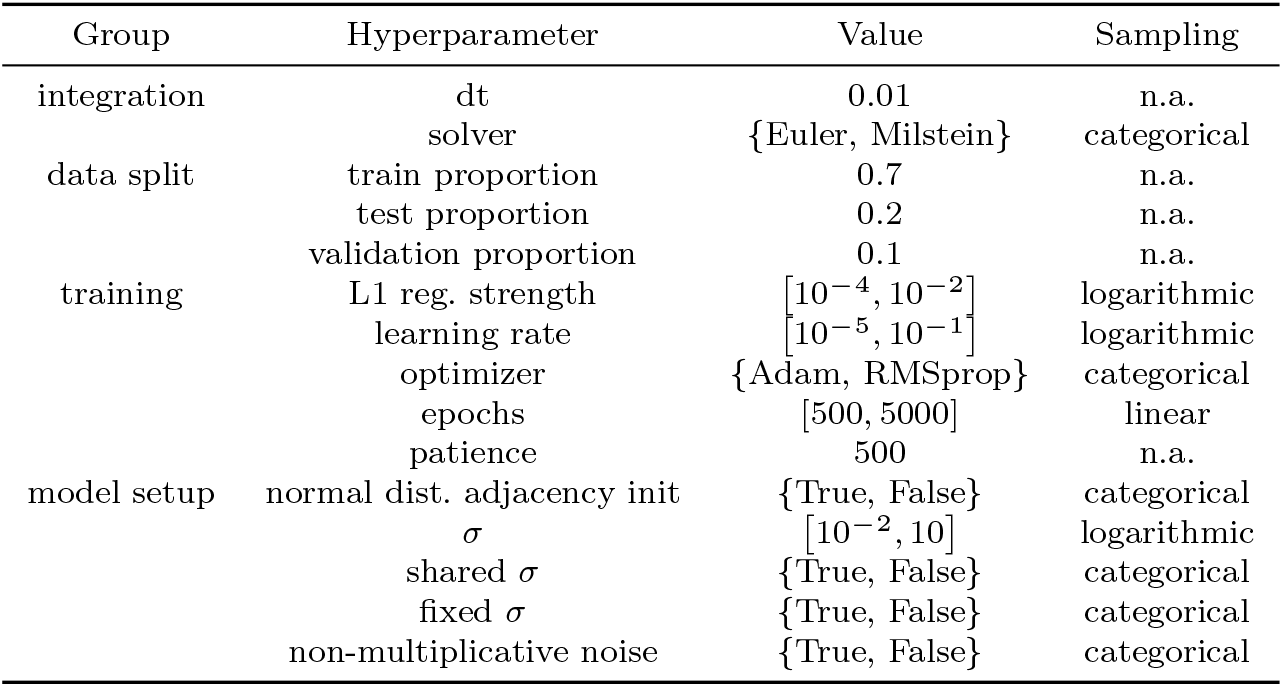
Model training hyperparameters for GRN benchmarking. Square brackets indicate a range of tested values, either sampled on a linear or logarithmic scale, and curly brackets indicate a categorical choice.

We used GENIE3 [26] and GRNBoost2 [27], among the best-performing methods in the BEELINE benchmark, especially for curated networks [24], as baseline methods for performance comparison. Specifically, we used the implementation of both methods in arboreto (v0.1.6) [27] and performed GRN inference for all 40 synthetic datasets (four curated networks, with ten datasets each).

To give a reference point for chance-level performance, we additionally built a random edge predictor for each dataset. For a dataset with *g* genes, we considered all *g*^2^ possible directed regulator–target pairs and sampled *e* of them uniformly at random without replacement, where *e* is the number of edges in the ground-truth network. Each selected edge received a distinct random rank as its score. All remaining pairs received the lowest score and were treated as tied. The random predictor therefore outputs a network of the same size as the ground truth, with its edges in random order. We repeated this procedure ten times per curated network.

For all three methods and the random estimator, we evaluated the performance based on the area under the receiver operating characteristic curve (AUROC), average precision (AP), and early precision (EP), where early precision is defined as the precision for the *e* highest-scoring edges, where *e* is the number of edges in the ground-truth network. For bowside, edge predictions were scored by absolute weight in the adjacency matrix. For GENIE3 and GRNBoost2, predicted edge importance was used as the score.

### Benchmarking trajectory inference

We used the same synthetic datasets as for GRN inference benchmarking to evaluate bowside’s cell fate prediction performance. Trajectories and cell fates were inferred using models trained with optimal hyperparameters (see Benchmarking GRN inference). The optimal number of trajectories was inferred automatically based on silhouette coefficients, with a minimum of two and a maximum of ten trajectories.

Palantir (v1.4.1) [6] and CellRank (v2.0.7) [28] were used as baseline models for performance comparison. For Palantir, diffusion maps were computed based on gene expression with palantir.utils.run diffusion maps with five diffusion components and 30 nearest neighbors. The multi-scale space was determined with palantir.utils.determine multiscale space. We chose the cell with the lowest overall pseudotime as the start cell, selected the cells with the highest trajectory-specific pseudotimes as the respective terminal states, and ran Palantir with 30 nearest neighbors and 500 waypoints. The terminal cells were explicitly specified because the automatic inference did not always find terminal states. This gives Palantir a slight advantage over the other methods, which did not receive any information about the expected number of trajectories or their corresponding terminal states.

For CellRank, we used two distinct kernels: a similarity-based kernel using the nearest-neighbor connectivity, and a pseudotime-based kernel. For each kernel, we fitted a GPCCA estimator [29], automatically selecting the number of macrostates between 3 and 20. The estimator was used to predict terminal states and fate probabilities.

We created ground-truth cell fate labels based on cell-level, trajectory-specific pseudotimes. Because each cell had an associated pseudotime value for only one of the two to five trajectories (depending on the curated model), cells were assigned to the corresponding trajectory. The caveat of this approach is that early, uncommitted cells also receive a single, fixed fate rather than similar probabilities for different fates. However, given the data, it is the most valid ground truth.

Based on these labels, we computed five performance measures: adjusted Rand index (ARI), adjusted mutual information (AMI), F1 score, balanced accuracy, and the area under the receiver operating characteristic (ROC) curve (AUROC). ARI and AMI compare the similarity of true and predicted label partitions and do not require a direct correspondence between true and predicted labels. In contrast, F1 score, balanced accuracy, and AUROC require matched predictions to measure classification accuracy. Because the expected number of trajectories and corresponding endpoints was fixed only for Palantir, CellRank and bowside did not necessarily predict the right number of trajectories, and the order was arbitrary. To obtain the optimal mapping of predicted labels to the ground truth, we solved the linear sum assignment problem to maximize the mean probability for each true trajectory based on the predicted fate probabilities. To compute balanced accuracy, we treated surplus, unmatched predicted trajectories as errors. Similarly, we computed the F1 score only for labels present in the ground truth, and treated additional labels as false or true negatives, depending on the respective ground truth. As the final F1 score, we report the unweighted mean of the per-class scores (macro averaging). For the AUROC computation, cells with an unmatched label received a probability of 0 for all ground-truth classes. The unweighted average of the one-vs-rest AUROC values for each true label against the rest, based on predicted fate probabilities, is reported as the final score.

As an additional random baseline, we permuted the true labels and assigned one-hot probabilities to maintain the true class proportions. This was done once per synthetic dataset, i.e., ten times for each of the four curated networks. We then computed the performance measures as described above.

## Acknowledgments

The authors thank the International Max Planck Research School for Intelligent Systems (IMPRS-IS) for supporting J.T.S. and M.Z. We thank the members of the Claassen group for valuable discussions and constructive feedback.

## Declarations

### Ethics approval and consent to participate

Not applicable.

### Consent for publication

Not applicable.

### Data availability

No new raw datasets were generated for this study. Processed data for reproducing the analyses are available on Zenodo at https://doi.org/10.5281/zenodo.23556963.

### Materials availability

Not applicable.

### Code availability

The bowside source code is available on GitHub at https://github.com/janschleicher/bowside, and a packaged version is available via the Python Package Index (https://pypi.org/project/bowside/). Code for reproducing the analyses and figures is available on GitHub at https://github.com/janschleicher/bowside_projects.

### Author contribution

*Jan T. Schleicher:* Conceptualization, Formal analysis, Investigation, Methodology, Software, Visualization, Writing – original draft, Writing – review & editing

*Marcello Zago:* Conceptualization, Investigation, Methodology, Writing – review & editing

*Manfred Claassen:* Conceptualization, Funding acquisition, Project administration, Resources, Supervision, Writing – original draft, Writing – review & editing

## Appendix A Extended data

**Fig. A1.**
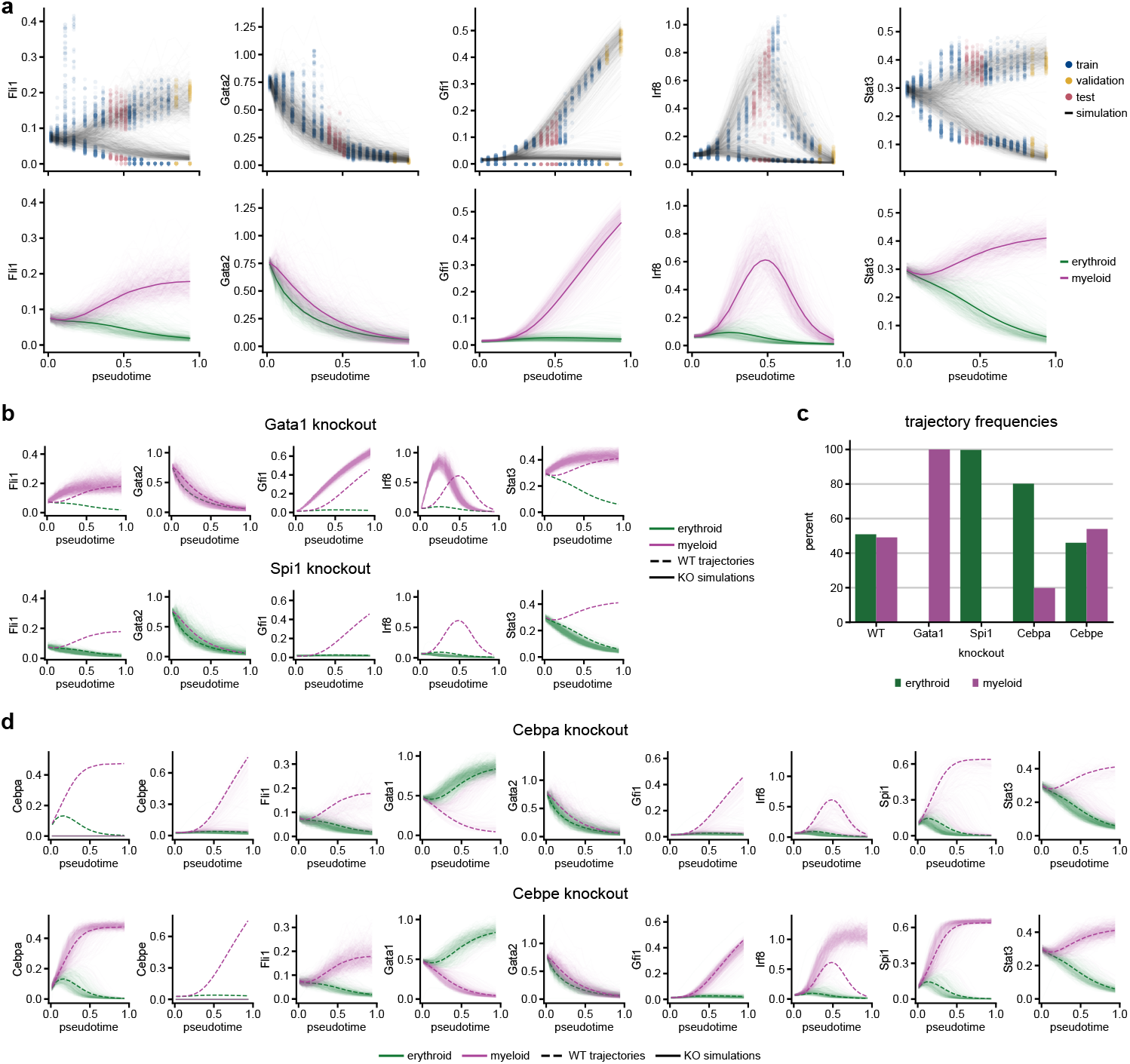
Extended data for. **Figure 1**. (a-b) Output of bowside for the remaining five genes not shown in Figure 1. (a) Simulations (top) and inferred trajectories (bottom) based on the trained model recapitulate branching gene expression. (b) Simulations of *Gata1* and *Spi1* knockouts. (c) Frequencies of the erythroid and myeloid trajectories for the wild-type and for knockouts of *Gata1*, *Spi1*, *Cebpa*, or *Cebpe* as the percentage of simulations predicted to belong to the respective trajectory. For the wild-type, simulations were assigned to a trajectory based on their cluster label. For the knockouts, the trajectory of a simulation was classified with a time series of Gaussian mixture models based on the wild-type hierarchical clustering. (d) Simulations of *Cebpa* (top) and *Cebpe* (bottom) knockouts.

**Fig. A2.**
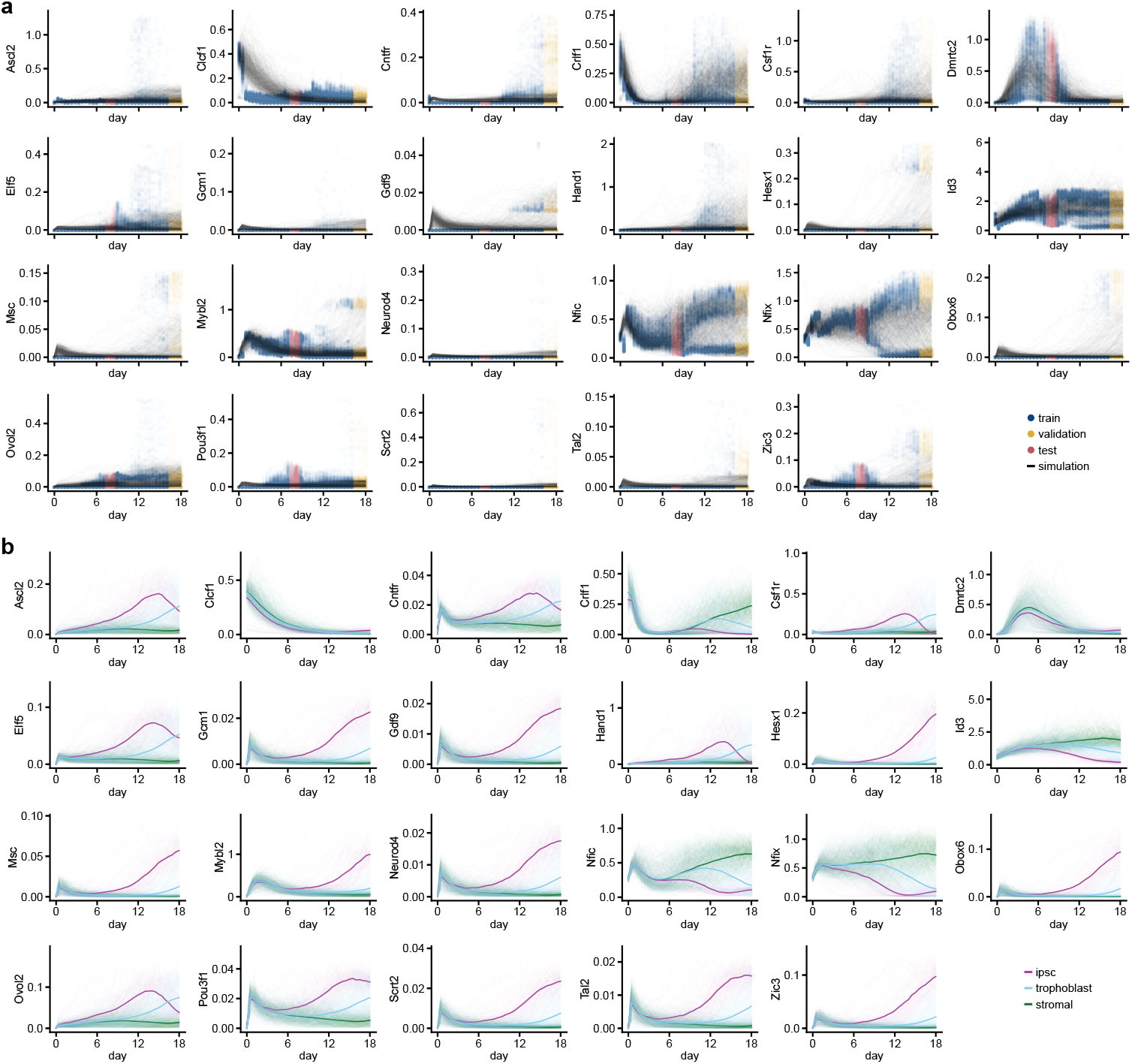
Extended data for. **Figure 2**. (a-b) Output of bowside for the remaining 23 genes not shown in Figure 2. Simulations (a) and inferred trajectories (b) based on the trained model.

**Fig. A3.**
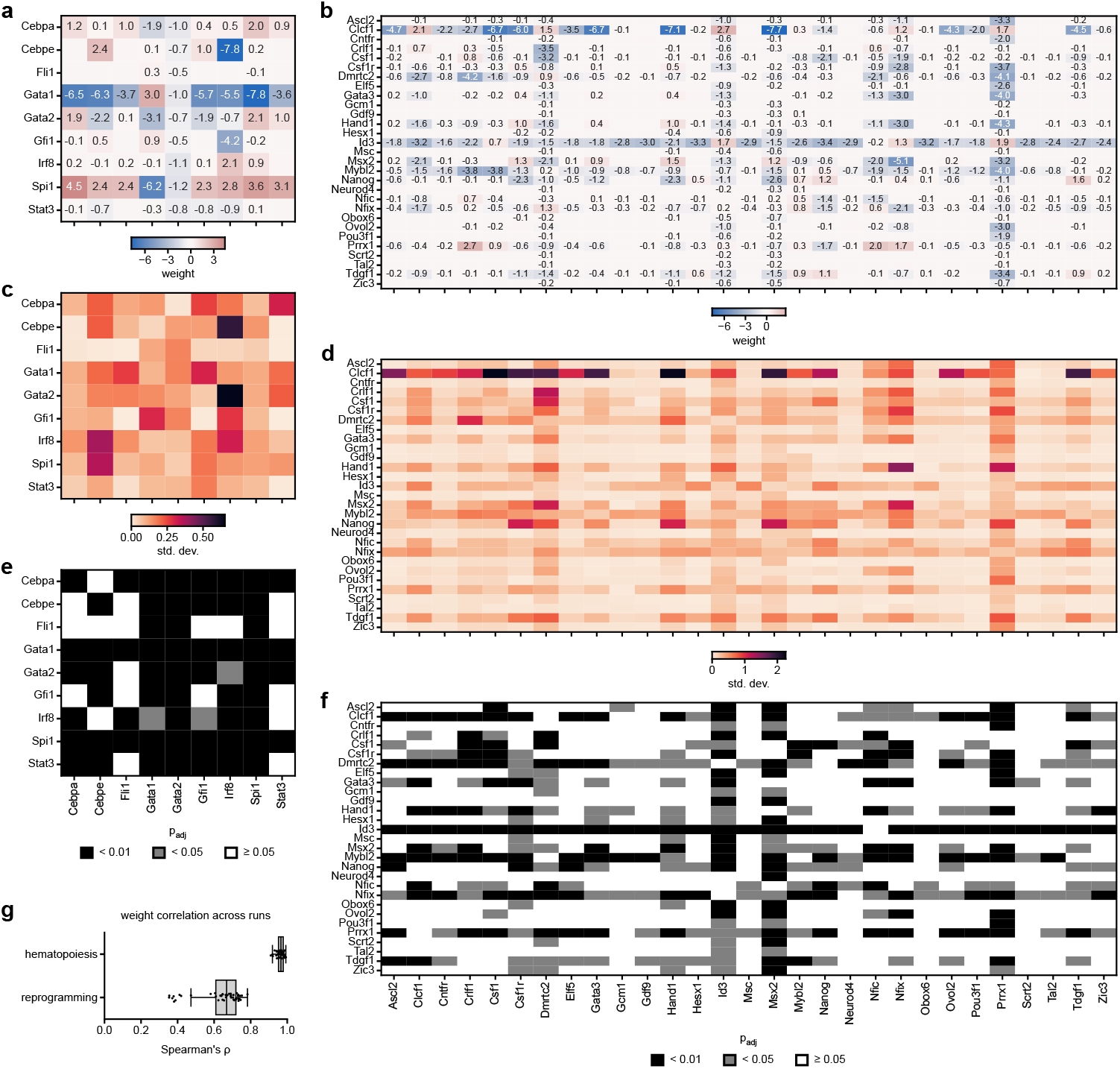
Stability of inferred GRN weights. Mean (a-b), standard deviation (c-d), and significance level (based on FDR-corrected Wilcoxon signed-rank test p-value, e-f) of adjacency matrix edge weights across ten independent model fitting repetitions with the same optimal hyperparameters for the hematopoiesis (a, c, e) and iPSC reprogramming (b, d, f) datasets, respectively. (g) Boxplot of Spearman’s rank correlation values for all pairwise comparisons between the inferred edge weights across ten model fitting repetitions.

**Fig. A4.**
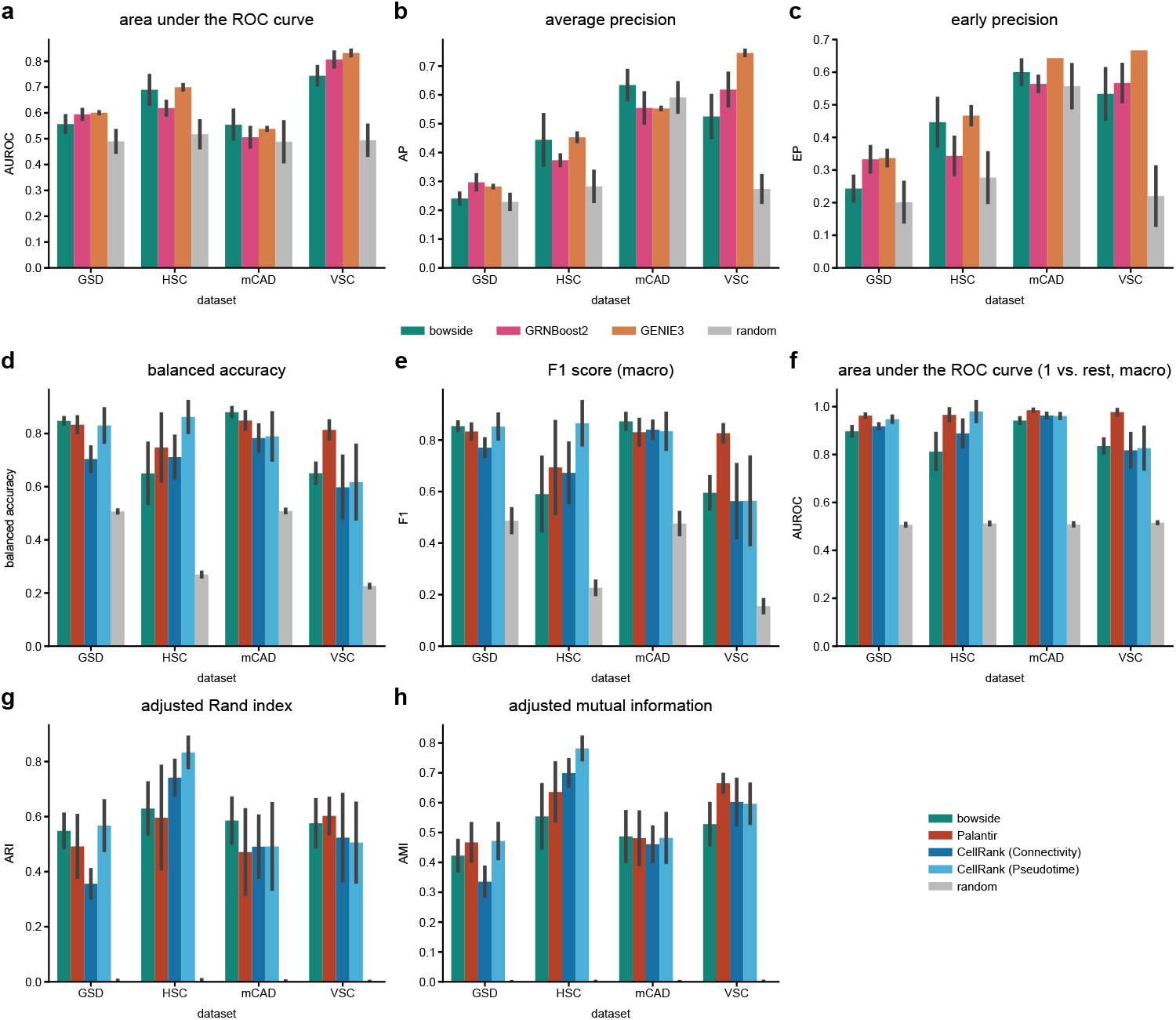
GRN and cell fate inference benchmark. (a-c) Comparison of GRN inference with bowside against two baseline methods and a random predictor on four synthetic curated datasets from BEELINE [24] in terms of (a) area under the receiver operating characteristic (ROC) curve (AUROC), (b) average precision (AP), and (c) early precision (EP). Mean of ten runs on ten distinct datasets for each curated model. Error bars show the standard deviation across runs. Random baseline: edges (equal to the number of ground-truth edges) sampled uniformly from all gene pairs and assigned random ranks; all other pairs scored zero; ten repetitions. (d-h) Comparison of cell fate inference with bowside against three baseline methods and a random predictor on four synthetic curated datasets from BEELINE [24] in terms of (d) balanced accuracy, (e) macro-averaged F1 score, (f) macro-averaged one-vs-rest area under the receiver operating characteristic (ROC) curve (AUROC), (g) adjusted Rand index (ARI), and (h) adjusted mutual information (AMI). Mean of ten runs on ten distinct datasets for each curated model. Error bars show the standard deviation across runs. Random baseline: random permutation of the true fate labels. One permutation per dataset.

